# Modelling the Putative Ancient Distribution of the Costal Rock Pool Mosquito *Aedes togoi*

**DOI:** 10.1101/2020.01.21.914838

**Authors:** Daniel A H Peach, Benjamin J Matthews

## Abstract

The coastal rock pool mosquito, *Aedes togoi*, is found in coastal east Asia in climates ranging from subtropical to subarctic. However, a disjunct population in the Pacific Northwest of North America has an ambiguous heritage. Two potential models explain the presence of *Ae. togoi* in North America: ancient Beringian dispersal or modern anthropogenic introduction. Genetic studies have thus far proved inconclusive. Here we described the putative ancient distribution of *Ae. togoi* habitat in east Asia and examined the climatic feasibility of a Beringian introduction into North America using modern distribution records and ecological niche modeling of bioclimatic data from the last interglacial period (∼120,000 BP), the last glacial maximum (∼21,000 BP), and the mid-Holocene (∼6000 BP). Our results suggest that suitable climatic conditions existed for *Ae. togoi* to arrive in North America through natural dispersal as well as to persist there until present times. Furthermore, we find that ancient distributions of suitable *Ae. togoi* habitat in east Asia may explain the genetic relationships between *Ae. togoi* populations identified in other studies. These findings indicate the utility of ecological niche modeling as a complementary tool for studying insect phylogeography.

## Introduction

The coastal rock pool mosquito, *Aedes* (*Tanakius*) *togoi* (Theobald), breeds in pools of brackish or salt-water above the high tide level on rocky shorelines and, sporadically, in containers of freshwater further inland (Petrishcheva 1948, Tanaka et al. 1979). *Ae. togoi* is a vector of the filarial parasite *Brugia malayi* (Tanaka et al. 1979, Hayashi 2011, Wada 2011), Japanese encephalitis (Petrishcheva 1948, Rosen et al. 1978, Rosen 1986) and potentially *Wuchereria bancrofti* and *Dirofilaria immitis* (Tanaka et al. 1979). *Ae. togoi* is found in coastal areas of east Asia, from subtropical to subarctic environments (Petrishcheva 1948, Tanaka et al. 1979, Sota et al. 2015), while a peripheral population of *Ae. togoi* also exists in the Pacific Northwest of North America along the coast of southern British Columbia, Canada (BC) and northern Washington, USA (Belton 1983, Darsie and Ward 2005, Peach 2018, Peach et al. 2019). The provenance of this population is unknown (Sota et al. 2015, Peach 2018), and two alternative hypotheses have been proposed to explain the presence of *Ae. togoi* in North America: invasion via anthropogenic dispersal and arrival via natural Beringian dispersal.

A trans-Beringian distribution is found in several insect species (Kavanaugh 1988), and many species of mosquito are Holarctic in distribution (Wood et al. 1979, Becker et al. 2010), but the restricted coastal habitat used by *Ae. togoi* may have limited it to the Pacific Rim. There are several reasons that the full distribution of *Ae. togoi* is not known. The remoteness and ruggedness of the region and Cold War politics have likely contributed to a dearth of sampling north of Japan. In North America, *Ae. togoi* do not tend to travel more than 20m from the shoreline (Trimble 1984) and thus trapping efforts outside of this narrow zone will not collect them. The rocky pools used by *Ae. togoi* is atypical mosquito habitat and is often difficult to access. Furthermore, larvae can remain submerged in detritus at the bottom of pools for extended periods of time (Sames et al. 2004). North American *Ae. togoi* have not been found breeding in containers (Trimble 1984, Sames et al. 2004), a common life-history trait of invasive mosquitoes (Hawley et al. 1987, Lounibos 2002). Conversely, there is some evidence for anthropogenic dispersal of *Ae. togoi* in parts of Asia (Sota et al. 2015) where it has been found on ships travelling between islands (Bohart 1956) and breeding in artifical containers (Tanaka et al. 1979) and the bilges of ships (Hsiao and Bohart 1946). One report suggests that certain populations of *Ae. togoi* in the Russian Far East can reside inland in freshwater containers associated with human habitation (Petrishcheva 1948). Furthermore, when suitable habitat is targeted for mosquito surveys *Ae. togoi* is regularly found (Sames et al. 2004), and it has been observed flying over tidewater to bite the occupants of boats at anchor in North America (Belton and Belton 1989).

*Ae. togoi* was initially described in Asia in 1907 (Theobald 1907) and first reported in North America from Victoria, BC in 1970 (Sollers-Riedel 1971). However, an undated specimen identified in the Canadian National Collection of Insect, Arachnids, and Nematodes in 1974 may have been collected as early as the 1940’s (Belton 1980). Furthermore, a 1919 record of *Aedes* (*Ochlerotatus*) *dorsalis* larvae from the supralittoral coastal rock pools at Caulfield Cove in North Vancouver, BC (Hearle 1926) has been proposed to be a mis-identification of *Ae. togoi* (Trimble and Wellington 1979). *Ae. dorsalis* is a floodwater mosquito that breeds in grassy, brackish tidal marshes (Wood et al. 1979, Belton 1983) while the permanent coastal rock pools at Caulfield Cove, at least currently (as of December 2019), host *Ae. togoi* and *Culiseta incidens* larvae (DP, Pers. Obs.). Thus, it is possible that *Ae. togoi* could have gone unnoticed in North America until sometime in the early-to-mid 20^th^ century.

Genetic evidence based on mitochondrial sequencing suggests that there are at least 4 distinct lineages of *Ae. togoi*: a lineage from temperate and subarctic regions of Japan, China, Taiwan, and other parts of Southeast Asia, a lineage from the subtropical islands of Japan, a lineage from subarctic Japan, and a lineage from Canada (Sota et al. 2015). If the *Ae. togoi* lineage in North America had arisen from natural divergence it is estimated to have diverged since some time in the Paleolithic era (Sota et al. 2015). This estimated divergence time spans several glacial cycles during which Beringian dispersal may have been possible. Alternatively, there may be unknown populations of *Ae. togoi* from which the North American population is derived (Sota et al. 2015). In addition to solving an entomological mystery, accurate accounting of *Ae. togoi* as invasive or indigenous has important implications. A change in status could alter *Ae. togoi*’s inclusion in analyses of invasion ecology/biology and help refine analyses of the attributes possessed by invasive mosquitoes. It could also properly inform legislative and other future action to combat invasive species and influence vector monitoring and control efforts. For example, we may not have to worry about the northward spread of *Ae. togoi* under climate change scenarios (e.g. Peach et al. 2019) if it is already indigenous to such areas.

Species distribution modeling, a type of environmental niche modeling, is a useful approach to estimate the suitability of habitat for a species in geographic areas it is not known to occupy or to estimate the changes in suitability of habitat for a species with environmental change over time (Wiens et al. 2009, Warren and Seifert 2011, Herrando-Moraira et al. 2019). Maximum entropy niche modelling (Maxent) is a commonly used approach in species distribution modelling (Rochlin et al. 2013, Melaun et al. 2015, Cunze et al. 2016, Wang et al. 2017, Peach et al. 2019, Rochlin 2019). Maxent is an effective open-source machine-learning algorithm that uses presence-only data to model habitat suitability (Elith et al. 2006, Phillips et al. 2006, 2017, Merow et al. 2013, West et al. 2016, Ashraf et al. 2017) and is frequently used to predict shifts in distribution of suitable habitat due to climate change (Rochlin et al. 2013, Wang et al. 2017, Peach et al. 2019) and putative distributions for species that become invasive in new regions (West et al. 2016, Peach et al. 2019, Rochlin 2019). Maxent modelling has also be applied to predict suitable habitat for a species under past climate conditions (Ye et al. 2014), including the paleodistribution of mosquitoes (Porretta et al. 2012). In the present study, we use available climate data to model the suitability of habitat for *Ae. togoi* along the Pacific Rim over the most recent glacial cycle to determine if suitable conditions existed for *Ae. togoi* to naturally disperse into and survive in North America. Moreover, we estimate the ancient distribution patterns of this mosquito in Asia and identify potential areas of allopatry that might be responsible for the distinct lineages found by Sota et al. (2015), as well as predict the current distributions of suitable *Ae. togoi* habitat that will help focus the search for as-yet unknown populations.

## Materials and methods

### Occurrence data

In the present study, we compiled 86 Asian records and 49 North American records of *Ae. togoi* within the study area (Figure 1, Supplementary Table 1). These records were taken from a combination of published literature (Petrishcheva 1948, Bohart 1956, Shestakov 1961, Lien 1962, Omori 1962, Kim and Seo 1968, Ramalingam 1969, Tanaka et al. 1975, 1979, Belton 1980, Sazonova and Smirnova 1986, Belton and Belton 1989, Sota 1994, Lee and Hong 1995, Sames et al. 2004, Stephen et al. 2006, Cheun et al. 2011, Sota et al. 2015, Petersen et al. 2017), personal collection records and observations (Gowlland Harbour, BC, Malcom Island, BC, Ucluelet, BC, Port Renfrew, BC, West Vancouver, BC, and Victoria, BC; DP, pers. obs.), and from museum specimens in the Beaty Biodiversity Museum in Vancouver, BC and the Royal British Columbia Museum in Victoria, BC. When identifiable place names were given rather than latitude and longitude, we derived associated coordinates from the centre of the location using Google Earth software (http://www.google.com/earth/download/ge).

**Figure 1:**
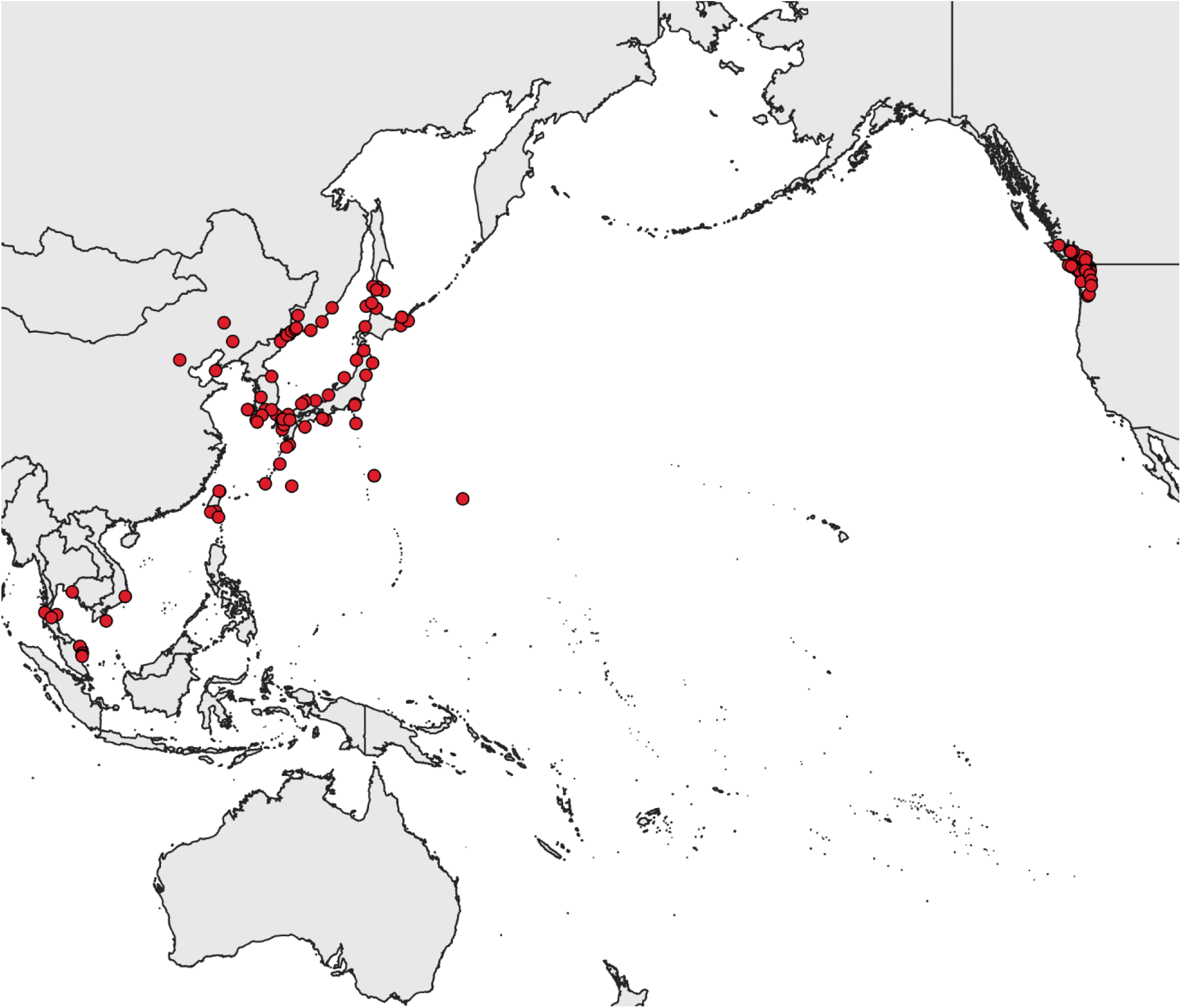
Worldwide distribution records of *Aedes togoi* used in this study (N = 135).

Some data points did not occur on pixels within bioclimatic data due to the irregular shape of coastline or the small size of islands. When this occurred, we relocated data points to the closest 1 km^2^ raster cell.

### Study area

We selected two study areas: one from Asia from within Latitude 0°N to 49°N and Longitude 94°E to 157°E (boundaries indicated on Figure 3), roughly encompassing the known distribution of *Ae. togoi* in Asia, and one for the North Pacific from within Latitude 0°N to 80°N and Longitude 95°E across the Pacific to 100°W, encompassing all known *Ae. togoi* populations in Asia and North America and possible unknown populations in the North Pacific. We projected a model from the Asian study area (hereafter referred to as model 1), obtained using Asian *Ae. togoi* populations, onto the North Pacific study area, following methodology for projecting habitat of invasive species into new areas (Cunze et al. 2018). We also generated results for the North Pacific study area using all known *Ae. togoi* populations (hereafter referred to as model 2), as if it were indigenous.

**Figure 2:**
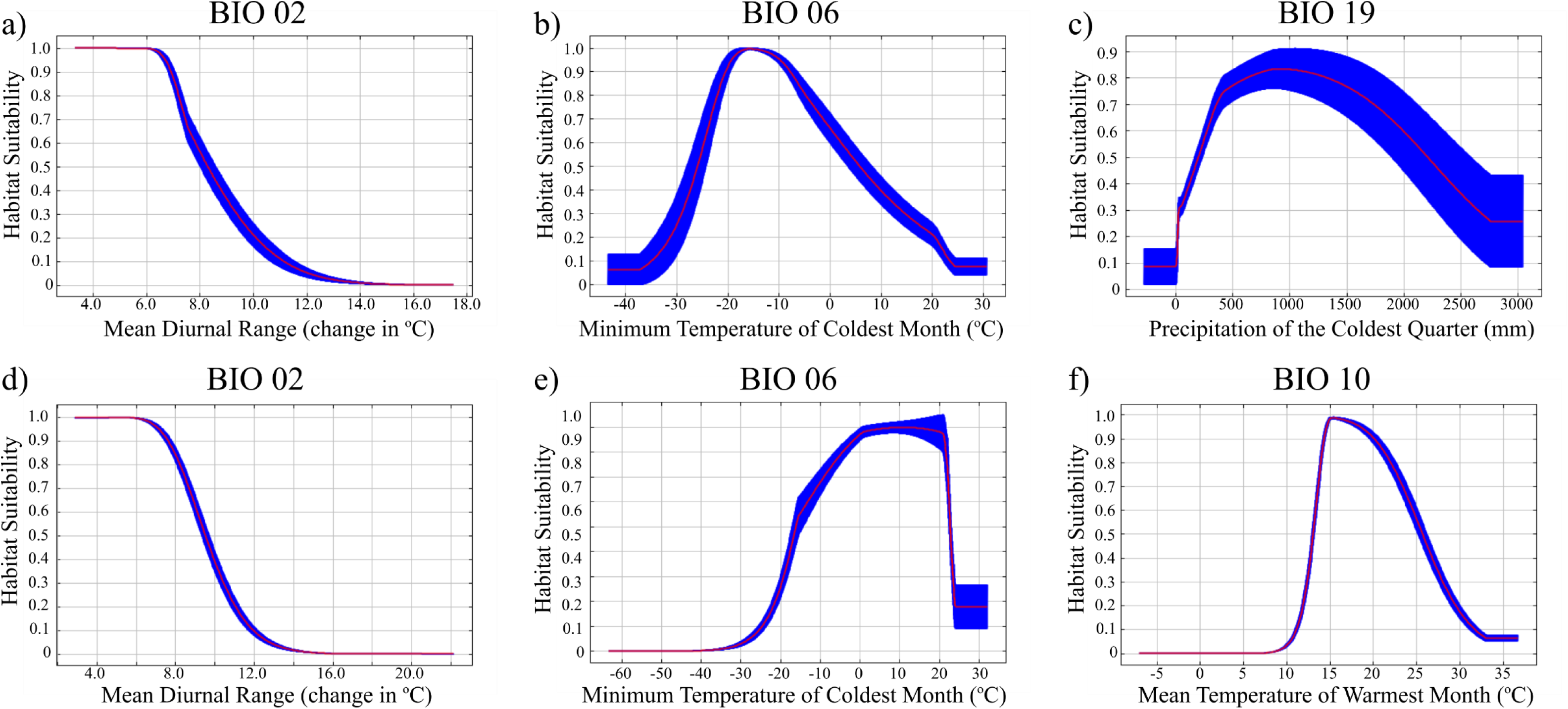
Relationship (mean ±SD) between environmental variables and Ae. togoi habitat suitability for model 1 (a,b,c) and model 2 (d,e,f).

**Figure 3:**
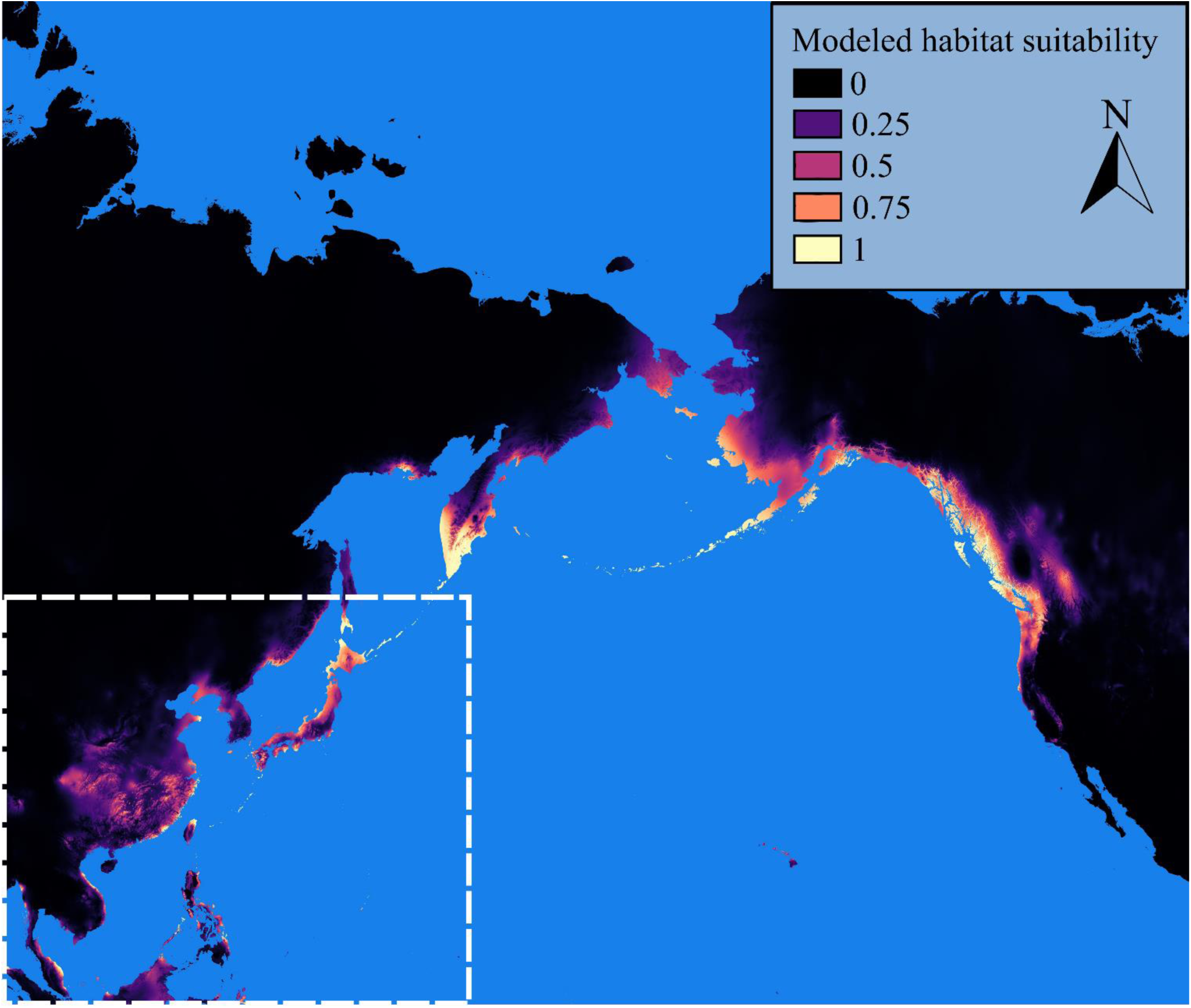
Current distribution of potential *Ae. togoi* habitat predicted by model 1, with hash marks indicating the Asian study area used to create model 1.

### Environmental variables

We downloaded 19 bioclimatic variables (Table 1) based on current climatic conditions at a scale of approximately 1 km^2^ (Hijmans et al. 2005) from Worldclim (http://worldclim.org-datasetversionv1.4, bioclimatic variables at 30s resolution). We also downloaded corresponding sets of variables based on climate conditions from: 1) the last interglacial period (∼120,000-140,000 BP) at a scale of 1 km^2^ (Otto-Bliesner et al. 2009), 2) the last-glacial maximum (∼22,000 BP) at a scale of 5 km^2^ (Gent et al. 2011), and 3) mid-Holocene period (∼6000 BP) at a scale of 1 km^2^ (Gent et al. 2011). We used these datasets to project our models onto past environments. We chose input bioclimatic variables based on possession of a weak intercorrelation with other variables, defined as a Pearson correlation <|0.8| calculated with ENMTools version 1.4.4 (Warren et al. 2010) (Supplementary Table 2, 3). When variables were highly correlated, we prioritized parameters for winter temperature due to relevance for *Ae. togoi* overwintering strategy (Sota 1994) and their importance in previous studies (Peach et al. 2019), parameters for precipitation due to reports of larval habitat drying out (Wada et al. 1993), and parameters for temperature of the wettest quarter due to their importance in previous studies (Peach et al. 2019). We calculated an Akaike information criterion (AICc) using ENMTools version 1.4.4 (Warren et al. 2010) and Maxent version 3.4.1 (Phillips et al. 2018) to compare models created with distinct environmental variables, model settings, and/or regularization parameters. We retained one best-performing model constructed from only Asian *Ae. togoi* observations (model 1) and all known *Ae. togoi* observations (model 2), each with different sets of 6 variables and clamping applied (Table 2).

**Table 1:** The Worldclim bioclimatic variables

| Code | Variable |
| --- | --- |
| BIO 01 | Annual Mean Temperature |
| BIO 02 | Mean Diurnal Range |
| BIO 03 | Isothermality |
| BIO 04 | Temperature Seasonality |
| BIO 05 | Maximum Temperature of the Warmest Month |
| BIO 06 | Minimum Temperature of the Coldest Month |
| BIO 07 | Temperature Annual Range |
| BIO 08 | Mean Temperature of the Wettest Quarter |
| BIO 09 | Mean Temperature of the Driest Quarter |
| BIO 10 | Mean Temperature of the Warmest Quarter |
| BIO 11 | Mean Temperature of the Coldest Quarter |
| BIO 12 | Annual Precipitation |
| BIO 13 | Precipitation of the Wettest Month |
| BIO 14 | Precipitation of the Driest Month |
| BIO 15 | Precipitation Seasonality |
| BIO 16 | Precipitation of the Wettest Quarter |
| BIO 17 | Precipitation of the Driest Quarter |
| BIO 18 | Precipitation of the Warmest Quarter |
| BIO 19 | Precipitation of the Coldest Quarter |

**Table 2:** The top candidate models based on AICc and AUC, with the final models constructed from Asian *Ae. togoi* records (model 1) and from all known *Ae. togoi* records (model 2) bolded.

| Occurrence Records | Variables | Parameters | Regularization | Features | AICc | AUC | SD |
| --- | --- | --- | --- | --- | --- | --- | --- |
| **Asia** | **Bio 02, 06, 10, 13, 18, 19** | **33** | **1** | **Without Linear** | **2600.3** | **0.912** | **0.031** |
| Asia | Bio 02, 06, 10, 13, 18, 19 | 25 | 2 | Without Linear | 2600.8 | 0.898 | 0.035 |
| Asia | Bio 02, 05, 06, 13, 18, 19 | 34 | 1 | Default | 2607.2 | 0.915 | 0.031 |
| **All** | **Bio 02, 06, 10, 14, 15, 16** | **37** | **1** | **Without Linear** | **4041.1** | **0.972** | **0.009** |
| All | Bio 02, 06, 10, 15, 16, 17 | 25 | 2 | Without Linear | 4042.2 | 0.971 | 0.01 |
| All | Bio 02, 06, 10, 15, 16, 17 | 41 | 1 | Without Quadratic | 4054.9 | 0.973 | 0.01 |

### Maxent

Maximum entropy niche modelling (Maxent) is a common approach for species habitat modelling (Melaun et al. 2015, Cunze et al. 2016, Peach et al. 2019, Rochlin 2019). Maxent is a machine-learning algorithm that models putative species distributions using presence-only data (Phillips et al. 2006, 2018). Maxent compares favourably to similar methods (Elith et al. 2006, Phillips et al. 2006, Padalia et al. 2014), even when minimal presence data is available (Elith et al. 2006, 2006). We used Maxent version 3.4.1 (Phillips et al. 2018) to model suitable habitat for *Ae. togoi*. We used default Maxent settings, default Maxent settings without linear features, and default Maxent settings without quadratic features, with 1000 replications, 20% of data points withheld for subsampling, clamping applied, and 10,000 background points. We analyzed results based on extensive presence of predicted suitable habitat on maps (original maps available to download at https://doi.org/10.5683/SP2/YPVTYT), with some areas of interest highlighted, and by the presence-only area under the curve (AUC_PO_) of the receiving operator characteristic (ROC). This integral gives a sum between 0 and 1 with a result >0.5 indicative of a model that is more accurate than random, >0.7 representing a useful model, >0.9 representing an excellent model, and a result of 1 indicating a perfect fit (Swets 1988). We compared the AUC_PO_ of our three best candidate modes as measured by AICc and selected the top performer across both metrics (Table 2). We also compared the relative contributions of the variables used by having Maxent randomly perform permutations of the value of each variable on presence and background data and then reassess the model. Decreases in AUC_PO_ were transformed to a percentage and used as a relative metric of variable importance. We built maps with QGIS version 3.4.3 (QGIS Development Team 2018), and downloaded spatial data on the extent of glaciation during the last glacial maximum (Ehlers et al. 2011) from the Collaborative Research Centre 806 Database (https://crc806db.uni-koeln.de/) to add glacial coverage layers where applicable.

## Results

### Model Selection and relevant variables

We retained six variables under default Maxent settings without linear features and with a regularization multiplier of 2 for model 1, and six variables under default Maxent settings without linear features and with a regularization multiplier of 1 for model 2 (Table 2). Both models used three shared variables: mean diurnal range, minimum temperature of the coldest month, and mean temperature of the warmest quarter. Model 1 has a mean AUC_PO_ of 0.912 (± 0.031) and model 2 has a mean AUC_PO_ of 0.972 (± 0.009), both representing models with excellent fits (Swets 1988).

For model 1, mean diurnal range, the minimum temperature of the coldest month, and precipitation of the coldest quarter were the relatively most important permutation variables (Table 3, Figure 2). For model 2, mean diurnal range, the minimum temperature of the coldest month, and mean temperature of the warmest quarter were the relatively most important permutation variables (Table 3, Figure 2).

**Table 3:** The relative permutation importance of the variables used in the *Ae. togoi* habitat models.

| Relative importance (%) to the Maxent models |  |  |
| --- | --- | --- |
| Variable | Model 1 | Model 2 |
| Mean diurnal range (BIO 02) | 46 | 24.3 |
| Minimum temperature of the coldest month (BIO 06) | 19.3 | 28.1 |
| Mean temperature of the warmest quarter (BIO 10) | 7.9 | 29.8 |
| Precipitation of the wettest month (BIO 13) | 9.2 | - |
| Precipitation of the driest month (BIO 14) | - | 8.9 |
| Precipitation seasonality (BIO 15) | - | 5.2 |
| Precipitation of the wettest quarter (BIO 16) | - | 3.6 |
| Precipitation of the warmest quarter (BIO 18) | 4.1 | - |
| Precipitation of the coldest quarter (BIO 19) | 13.4 | - |

### Current Conditions

Under current conditions our models predict possible *Ae. togoi* habitat on the North Coast of British Columbia, the Alexander Archipelago, Cook Inlet, and parts of the Aleutian Islands of Alaska, eastern Kamchatka, the Kurile Islands, the Philippines, and the Hawaiian Islands (Figures 3 & 4). Model 1 shows much more extensive suitable habitat for *Ae. togoi* species around most of the North Pacific, including areas not predicted by model 2 such as Taui Bay in the sea of Okhotsk, the northern Kurile Islands, most of Kamchatka, most of the Aleutian Islands, the Pribolif Islands, the coasts of the Bering Sea, and most of the coast of Alaska (Figure 3).

**Figure 4:**
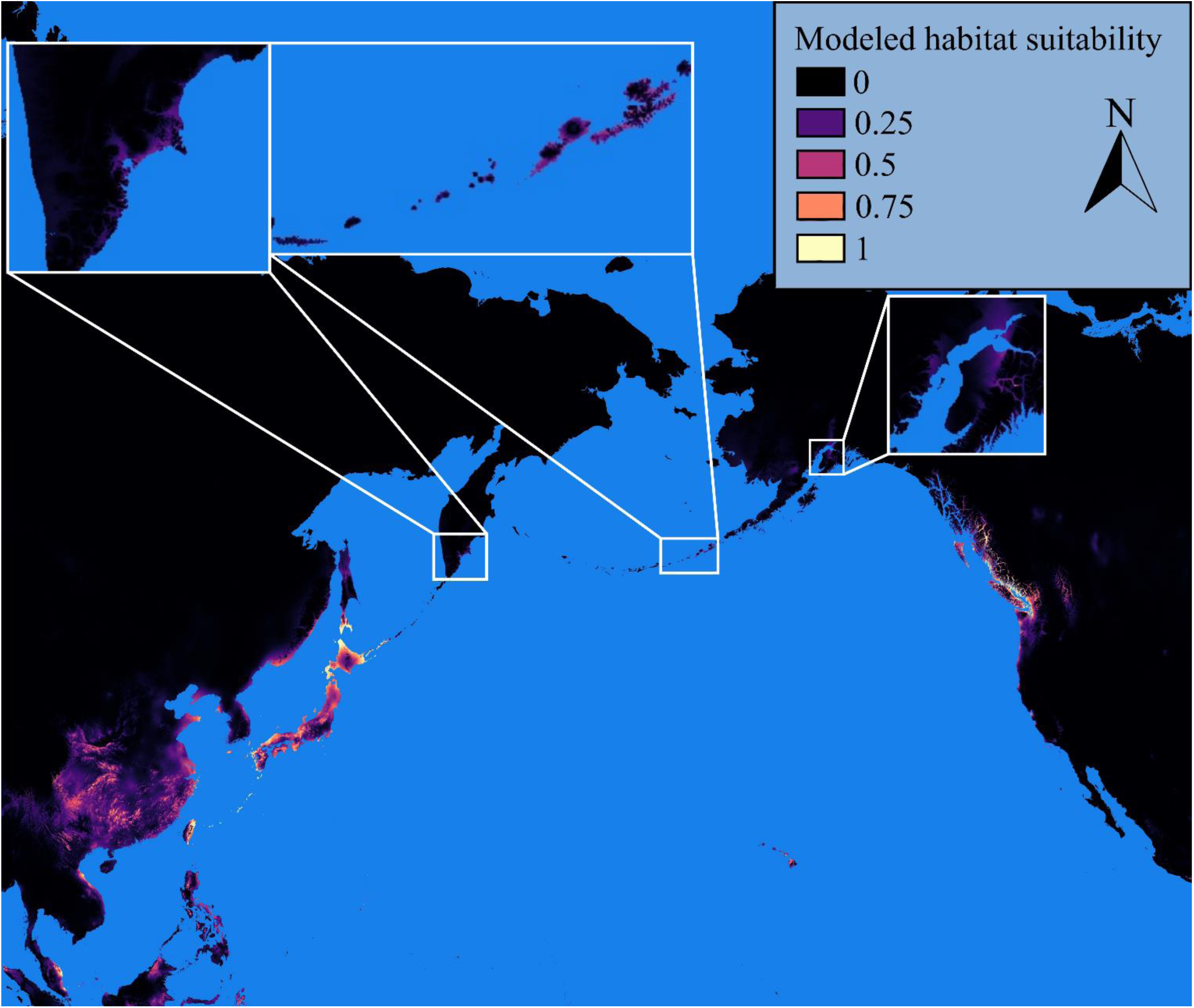
Current distribution of potential *Ae. togoi* habitat predicted by model 2.

### Last Interglacial Period

Our models both show a reduction of suitable habitat for *Ae. togoi* in mainland Asia during the last interglacial period (Figures 5 & 6); however, in general the predicted habitat for *Ae. togoi* during this time period are roughly similar to those predicted by our models for the current time period. One exception is that our model 2 predicts increased areas of suitable habitat in Kamchatka, the Aleutians, mainland Alaska, the Alexander Archipelago, and coastal British Columbia during the last interglacial period (Figure 6).

**Figure 5:**
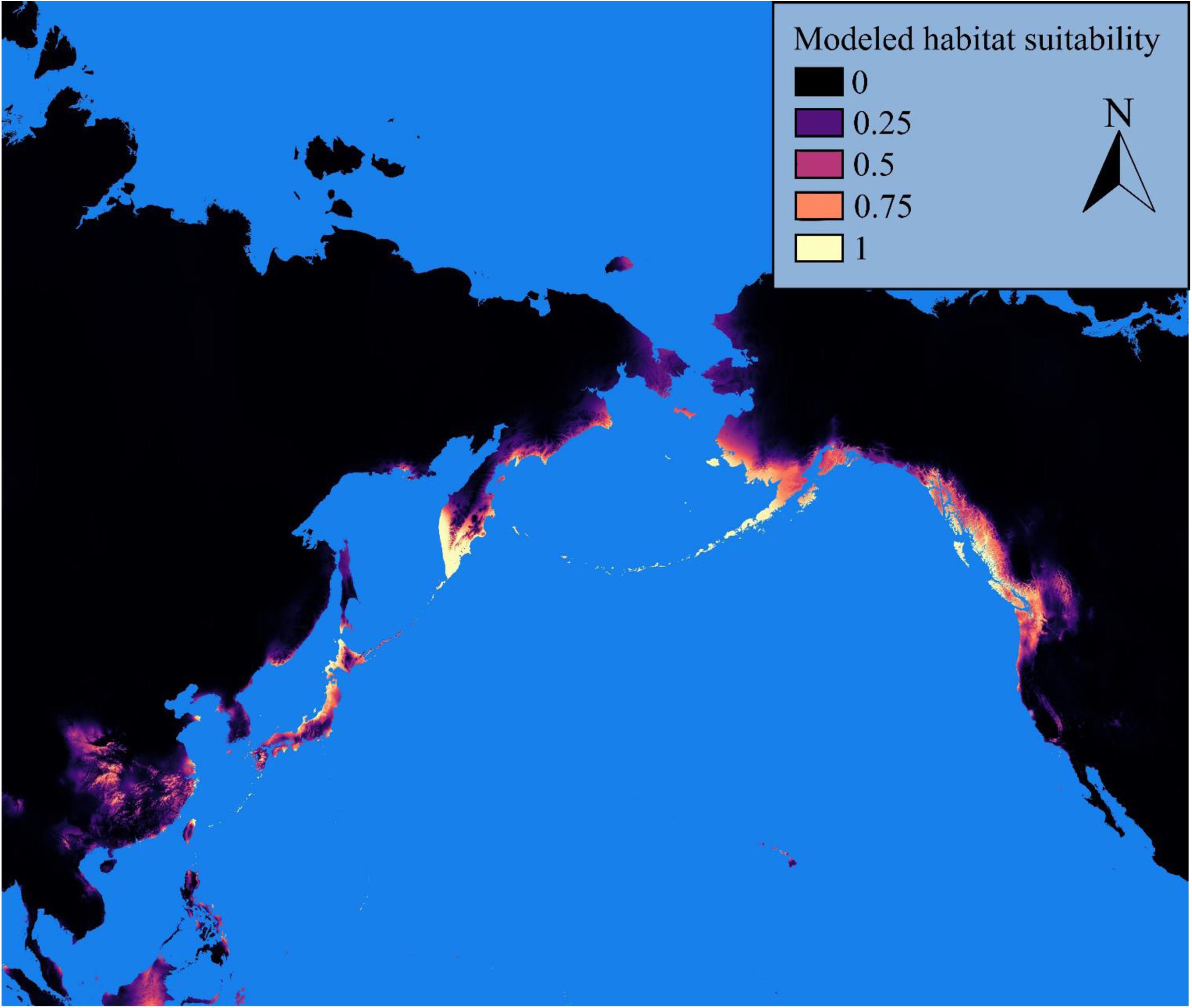
Paleodistribution of potential *Ae. togoi* habitat during the last interglacial period predicted by model 1.

**Figure 6:**
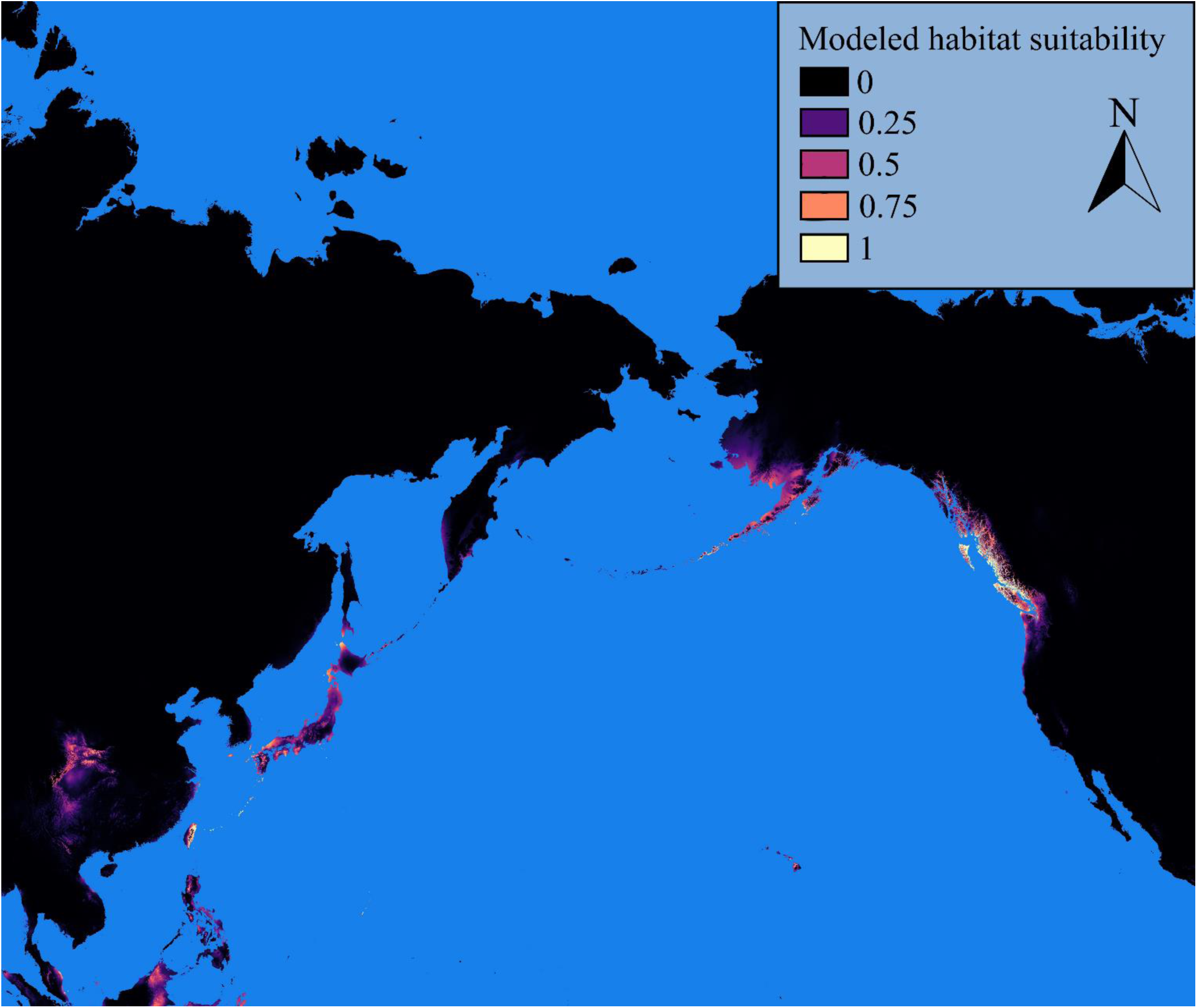
Paleodistribution of potential *Ae. togoi* habitat during the last interglacial period predicted by model 2.

### Last Glacial Maximum

Our model 1 predicts that ample habitat suitable for *Ae. togoi* was present along the coast of the Bering land bridge, the Aleutian Islands, and the coast of North America from Alaska to the Baja during the last glacial maximum (Figure 7). While much of this area was covered by ice sheets, our model predicts that patches of exposed rock pools along the coast could have provided suitable microhabitats for *Ae. togoi*. Moreover, ice-free glacial refugia such as portions of the Aleutian Islands, the Alexander Archipelago, Haida Gwaii, and northern Vancouver Island are predicted to be suitable habitat for *Ae. togoi*. Model 1 also suggests regions of suitable habitat for *Ae. togoi* in eastern Asian during this time period including Japan’s main islands, the Ryukyu archipelago, the Izu Archipelago, the Bonin Islands coastal China, parts of the Philippines, and along the coast of the South China sea and the islands it contained (Figure 7). Conversely, our model 2 predicts that suitable habitat for *Ae. togoi* during the last glacial maximum was restricted to a small region of coastal California, Baja California, and the Channel Islands of California (Figure 8). Model 2 suggests largely the same distribution of suitable habitat for *Ae. togoi* in eastern Asia during the last glacial maximum as model 1 (Figure 8).

**Figure 7:**
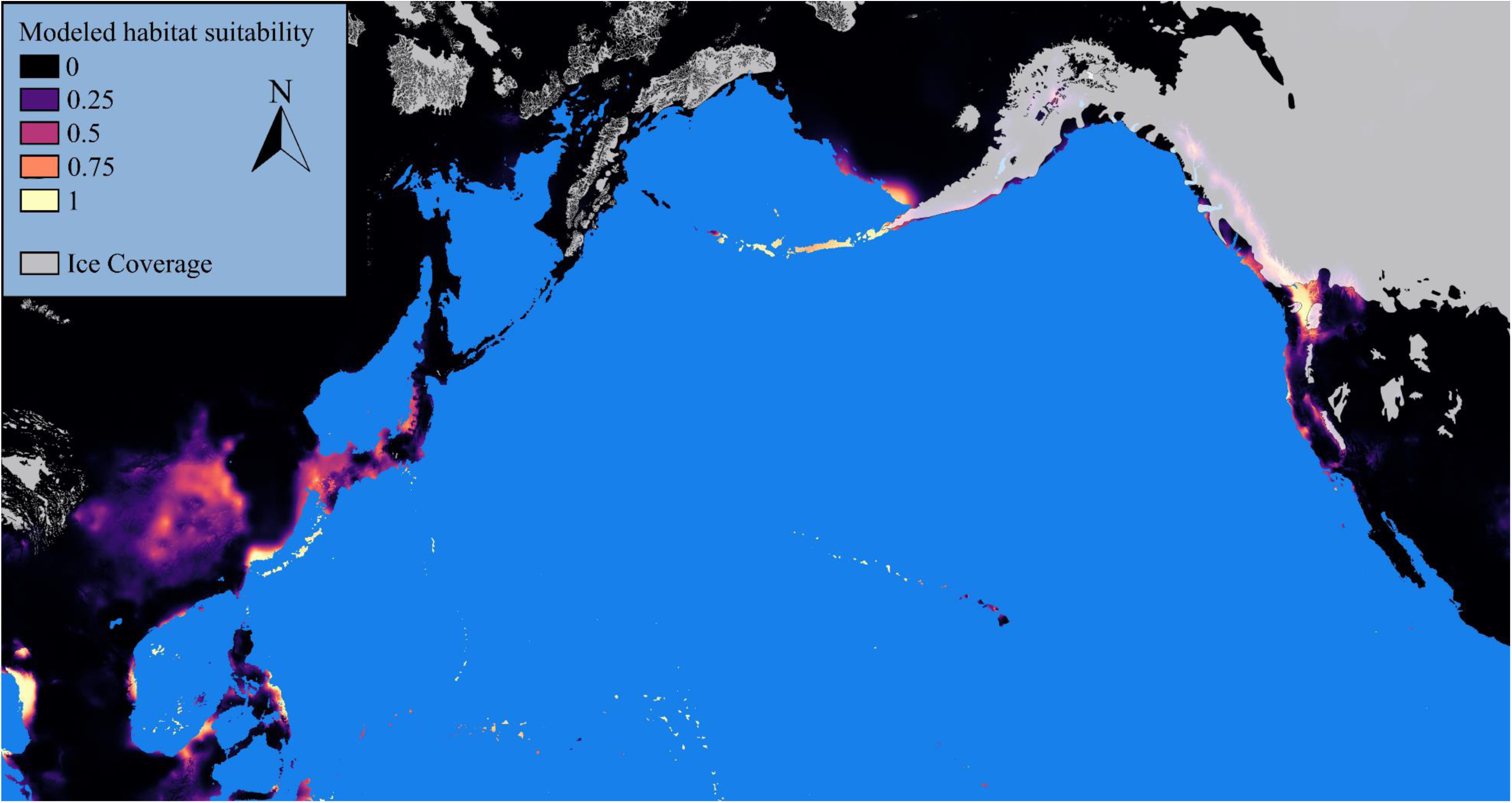
Paleodistribution of potential *Ae. togoi* habitat during the last glacial maximum predicted by model 1, with ice coverage.

**Figure 8:**
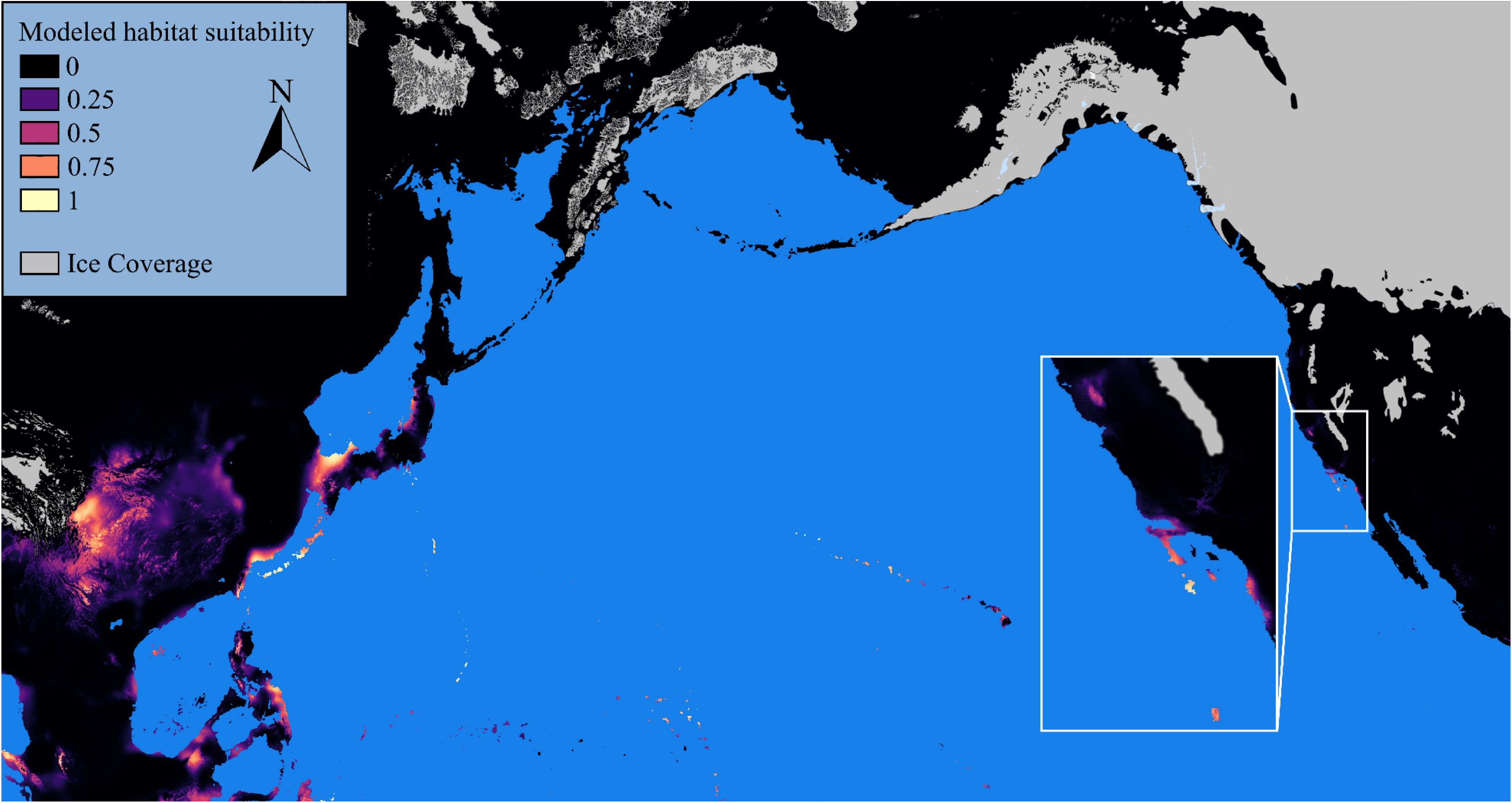
Paleodistribution of potential *Ae. togoi* habitat during the last glacial maximum predicted by model 2, with ice coverage.

### Mid-Holocene

Our models both predict that the distribution of *Ae. togoi* habitat in the mid-Holocene was roughly similar to the present-day distribution of *Ae. togoi* (Figures 9 & 10). Differences include model 1 predicting reduced *Ae. togoi* habitat particularly along the coast of the Gulf of Alaska and the coastlines of the Bering Sea (Figure 9), whereas model 2 predicts reduced habitat in Cook Inlet and the Alexander Archipelago (Figure 10).

**Figure 9:**
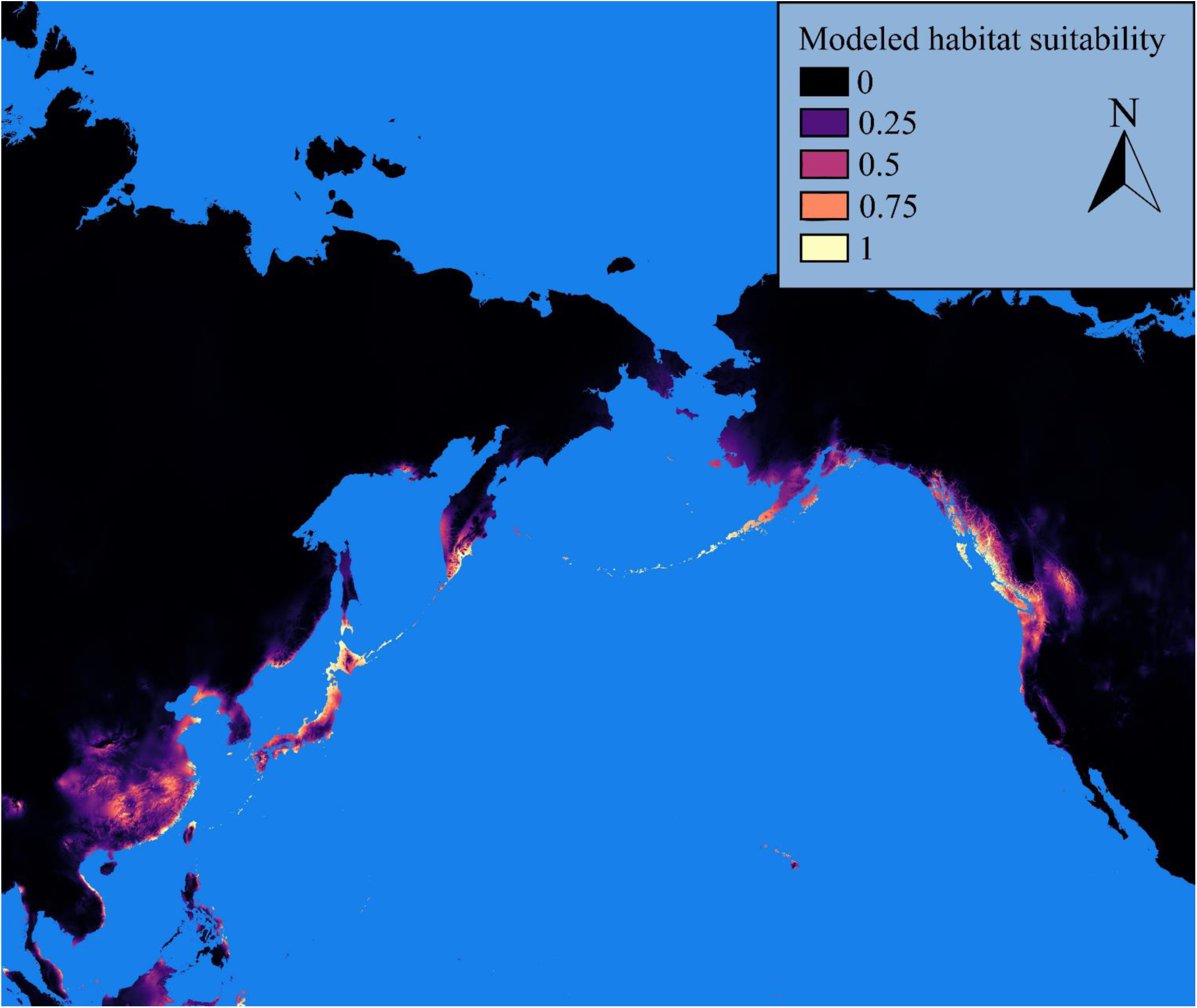
Paleodistribution of potential *Ae. togoi* habitat during the mid-Holocene period predicted by model 1.

**Figure 10:**
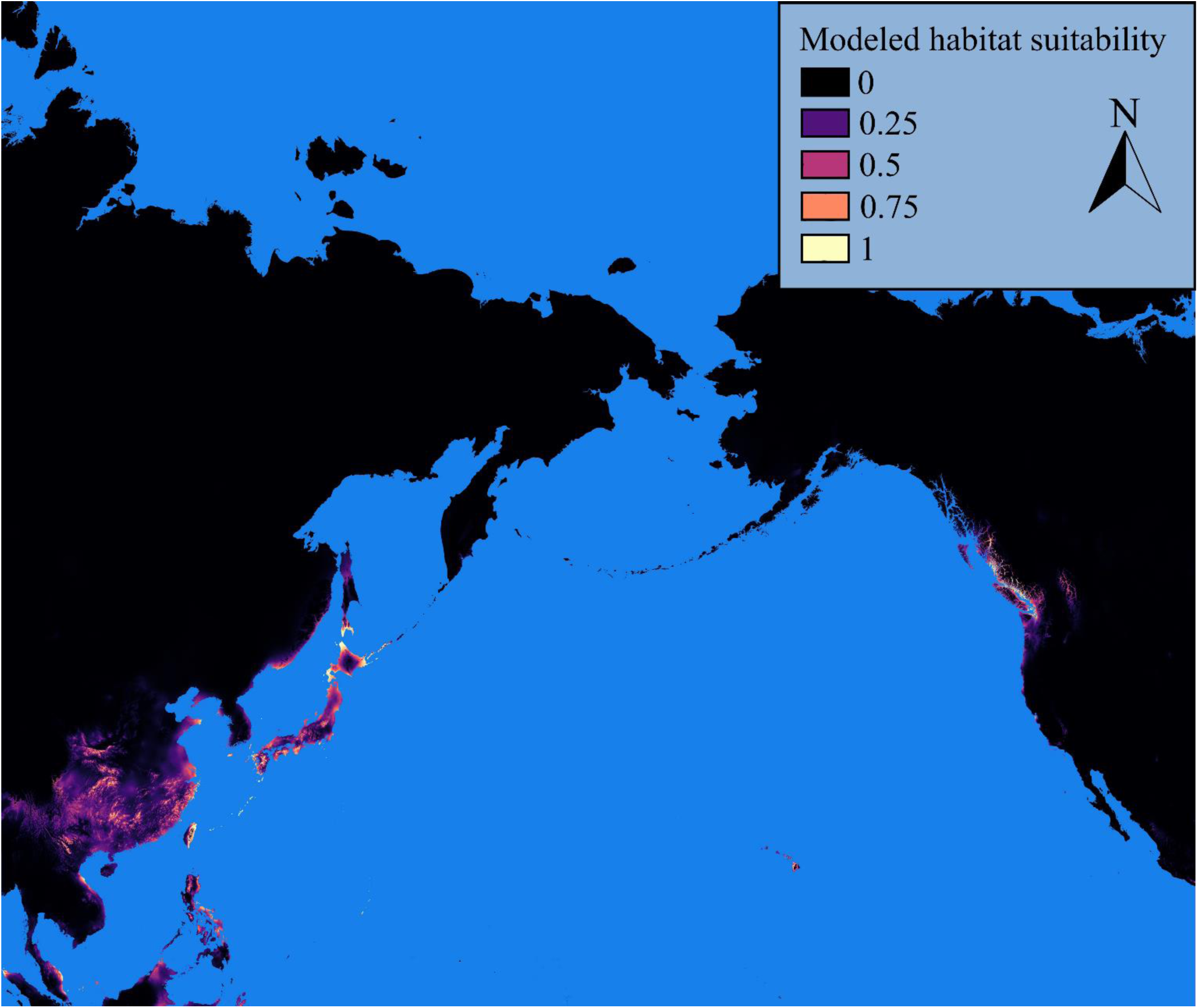
Paleodistribution of potential *Ae. togoi* habitat during the mid-Holocene period predicted by model 2.

## Discussion

Our models predict the presence of suitable climate for *Ae. togoi* around much of the North Pacific, including regions of North America and Far Eastern Russia without existing observations of *Ae. togoi*. These areas, including parts of Kamchatka, the Aleutian Islands, Cook Inlet, the Alexander Archipelago, and the north Coast of British Columbia, are remote and rugged terrain that has very likely been under-surveyed. Therefore, in agreement with Sota et al. (2015), we predict that undiscovered *Ae. togoi* populations may exist in the North Pacific.

Model 1, which uses only Asian observations of *Ae. togoi*, shows a much greater predicted distribution of *Ae. togoi* habitat than model 2, which additionally included North American observations. Projecting species habitat suitability into a novel range using only training data from the original range can lead to overestimation of projected suitable habitat due to local or regional differences in the ecological adaptations of sub-populations which, when taken in aggregate, can overestimate the species’ ecological breadth (Stockwell and Peterson 2002). Furthermore, *Ae. togoi* almost exclusively uses supralittoral rock pools as breeding habitat and, accordingly, results from both models should be interpreted to mean that only subsets of the environmentally suitable range that contain suitable breeding habitat are potential niches for *Ae. togoi*. Interestingly, both of our models predict suitable conditions for *Ae. togoi* in the Hawaiian Islands, as have previous studies (Peach et al. 2019), as well as other Pacific Islands such as Guam and the Northern Mariana Islands.

Both of our models predict that putative *Ae. togoi* habitat existed along the Pacific coast of North America during the last interglacial period, last glacial maximum, mid-Holocene, and present day. However, our model 2 predicts that suitable *Ae. togoi* habitat during the last glacial maximum was restricted to California, the Channel Islands of California, and Baja California. Whether or not *Ae. togoi* could have followed the shifting distribution of suitable environmental conditions to this area as North America cooled is unknown. While much of northern North America was covered by ice sheets during the last glacial maximum (Shafer et al. 2010, Ehlers et al. 2011), *Ae. togoi* could have persisted in one or more glacial refugia such as the Aleutians, the Alexander Archipelago, Haida Gwaii, northern Vancouver Island, or other cryptic refugia (Shafer et al. 2010). This continual presence of suitable *Ae. togoi* habitat in North America throughout a complete glacial cycle, along with suitable ancient habitat connecting North America to Asian *Ae. togoi* populations, implies that this species could have naturally dispersed into North America and survived long-term climatic cycles into the present. We note that presence-only models can be inaccurate when produced from data that suffers from sampling bias (Yackulic et al. 2013) that and our models may suffer from limitations in predicting paleodistributions of suitable *Ae. togoi* habitat due to a dearth of sampling records from remote and difficult to access regions in the northern coastal regions of *Ae. togoi’s* present range.

The present study suggests that *Ae. togoi* habitat has existed in much of coastal southeast Asia throughout the last glacial cycle and that its distribution may partially explain the relationships between geographically distinct populations described in other studies. The presence of suitable *Ae. togoi* habitat in the Ryukyu archipelago predicted by both of our models throughout all time frames examined supports, and may partially explain, the unique lineage of *Ae. togoi* populations described on these islands (Sota et al. 2015). The continuous presence of *Ae. togoi* on the Ryukyu Archipelago implied by our models, coupled with a barrier to gene flow, may have allowed for the evolution of an *Ae. togoi* sub-population with specific adaptations to these islands. Widespread and largely connected *Ae. togoi* habitat along the coasts of east and southeast Asia throughout the last glacial cycle may also explain the widespread distribution of the single *Ae. togoi* lineage found from subarctic to subtropical locations (Sota et al. 2015). We believe that the benefit of linking population genetic structure to geography across distinct evolutionary scenarios is a key benefit of including studies of paleodistribution from species distribution modelling techniques in phylogeography (Porretta et al. 2012).

Mean diurnal range and minimum temperature of the coldest month were important environmental variables in each of our models. In addition, precipitation during the coldest quarter was also critical in model 1, while mean temperature during the warmest quarter was important in model 2. There is evidence that, under certain conditions, temperature fluctuations (rather than absolute temperature) can have negative effects on the development rate and survival of some mosquitoes (Lyons et al. 2013). With this in mind, the importance of mean diurnal range may reflect a biological reliance on relatively stable environmental temperature conditions during *Ae. togoi* larval development. Alternatively, the importance of mean diurnal temperature range in our models may simply reflect that *Ae. togoi*’s breeding habitat is close to the ocean and subject to its stabilizing effect on temperature. The minimum temperature of the coldest month is an important environmental variable for the habitat of other *Aedes* spp. (Melaun et al. 2015, Lubinda et al. 2019), and another measure of cold tolerance, the mean temperature of the coldest quarter, was found to be important for *Ae. togoi* habitat suitability in North America (Peach et al. 2019). Such parameters are important determinants of overwintering strategy (Sota 1994) and overwinter survival of some mosquitoes (Kaufman and Fonseca 2014). Insulation of mosquito breeding habitat by snow may help to protect eggs and larvae from the effects of low air temperature (Hanson and Craig 1995) and could be responsible for the importance of precipitation during the coldest quarter as a variable in model 1. *Ae. togoi* breeding habitat does occasionally dry out (Wada et al. 1993), and it could also be that this variable simply reflects the danger of these pools drying out when the mosquitoes are restricted to the egg or larval stage and at their most vulnerable to desiccation. The importance of mean temperature of the warmest month on *Ae. togoi* habitat suitability in model 2 may reflect the well-known importance of temperature in mosquito development times andrvival (Alto and Juliano 2001, Reinhold et al. 2018). This variable is also important in for the habitat suitability of other *Aedes* spp. (Cunze et al. 2016). Despite it’s importance in other studies (Peach et al. 2019), the mean temperature of the wettest quarter was not part of our best-fitting models. This could be due to the different geographic areas used in our models.

Trade from Japan has been suggested as a potential vehicle for the introduction of *Ae. togoi* into North America (Belton 1983, Sota et al. 2015), but there was Russian activity related to colonization, the fur trade, whaling, collection and shipping of salt, and other commercial activities on the Pacific Coast of North America as far back as the 18^th^ century (Gibson 1980, Lightfoot 2003). This exchange between Northeast Russia and Pacific North America by the Russian American Company and its predecessors, involving Russian ports in the Sea of Okhotsk, Amur region, and Kamchatka (Gibson 1980), could also have been responsible for the introduction of *Ae. togoi* to North America. Interestingly, the ships used to transport and supply Canadian soldiers and the Royal Northwest Mounted Police, deployed to combat Bolsheviks in Vladivostok and Siberia as part of the Canadian Siberian Expeditionary Force (Isitt 2006), could also have been responsible. The timing of these forces’ return to Canada in 1919 (Isitt 2006) lines up conspicuously well if Hearle’s 1919 record of *Ae. dorsalis* from coastal rock pools in Caulfield Cove, West Vancouver (Hearle 1926), was mistaken from *Ae. togoi* (Trimble and Wellington 1979).

Ultimately, our modeling is based on climate data alone and is subject to the realities of limited sampling and, thus, cannot definitely prove or disprove either an anthropogenic introduction or Beringian dispersal for the presence of *Ae. togoi* in North America. However, models such as these can be used to inform survey efforts aimed at detection of additional populations of species of interest (Law et al. 2017, Rhoden et al. 2017). We propose that future survey efforts for *Ae. togoi* should be directed at predicted areas of suitable habitat in Kamchatka, the Aleutian Islands (particularly Umnak and Unalaska), Cook Inlet, the Alexander Archipelago, and the North Coast of British Columbia. We also note a need for *Ae. togoi* surveillance across its novel predicted range in more heavily-surveyed areas, including the Hawaiian Islands and the Mariana Archipelago. To further investigate the origins of *Ae. togoi* in North America, we propose a combination of survey efforts and population genetic analyses based on mitochondrial and nuclear genome sequencing (Lee et al. 2019).

## Supporting information

Unix Code For Model Runs

Unix Code For Model Selection

## Acknowledgements

We thank Karen Needham of the UBC Beaty Biodiversity Museum and Claudia Copley of the Royal BC Museum for access to specimens. We additionally thank Alistair Blachford of the Zoology Computing Unit, UBC for technical assistance and L. Melissa Guzman for helpful discussions.

